# Wikidata: A platform for data integration and dissemination for the life sciences and beyond

**DOI:** 10.1101/031971

**Authors:** Elvira Mitraka, Andra Waagmeester, Sebastian Burgstaller-Muehlbacher, Lynn M. Schriml, Andrew I. Su, Benjamin M. Good

## Abstract

Wikidata is an open, Semantic Web-compatible database that anyone can edit. This ‘data commons’ provides structured data for Wikipedia articles and other applications. Every article on Wikipedia has a hyperlink to an editable item in this database. This unique connection to the world’s largest community of volunteer knowledge editors could help make Wikidata a key hub within the greater Semantic Web. The life sciences, as ever, faces crucial challenges in disseminating and integrating knowledge. Our group is addressing these issues by populating Wikidata with the seeds of a foundational semantic network linking genes, drugs and diseases. Using this content, we are enhancing Wikipedia articles to both increase their quality and recruit human editors to expand and improve the underlying data. We encourage the community to join us as we collaboratively create what can become the most used and most central semantic data resource for the life sciences and beyond.

## 1 Stone Data Soup

In the Stone Soup folktale [1], a group of hungry travelers arrive in a village with its inhabitants unwilling to share their food. With a kettle of water and a stone the travelers manage to touch the curiosity of the villagers. The curiosity finally spawns a collaborative effort to make a great soup. This story is nowadays used to express the power of crowdsourcing and collaborative projects [2], such as Wikipedia, where many individuals each make small contributions but collectively produce something larger than the sum of its parts. Wikidata extends this collaborative model to the Web of data [3]. In this article we will describe Wikidata and the ways that this open public platform can take a central role in data sharing and management for the life science community.

## 2 Wikidata and Wikipedia

Wikipedia is among the most visited sites on the Internet. Articles about medical topics were viewed more than 4.88 billion times in 2013, a number on par with http://nih.gov and significantly greater than WebMD [4]. This incredibly important resource, created through volunteer labor, is now tightly coupled to Wikidata – an open, Semantic Web-compatible database that anyone can edit [3]. Wikipedia infoboxes – the tables of data often appearing on the right side of articles – can now render content stored in Wikidata and each Wikipedia article now has a direct link to the corresponding Wikidata item, thus encouraging the collaborative editing of the data (Fig. 1).

**Fig. 1.**
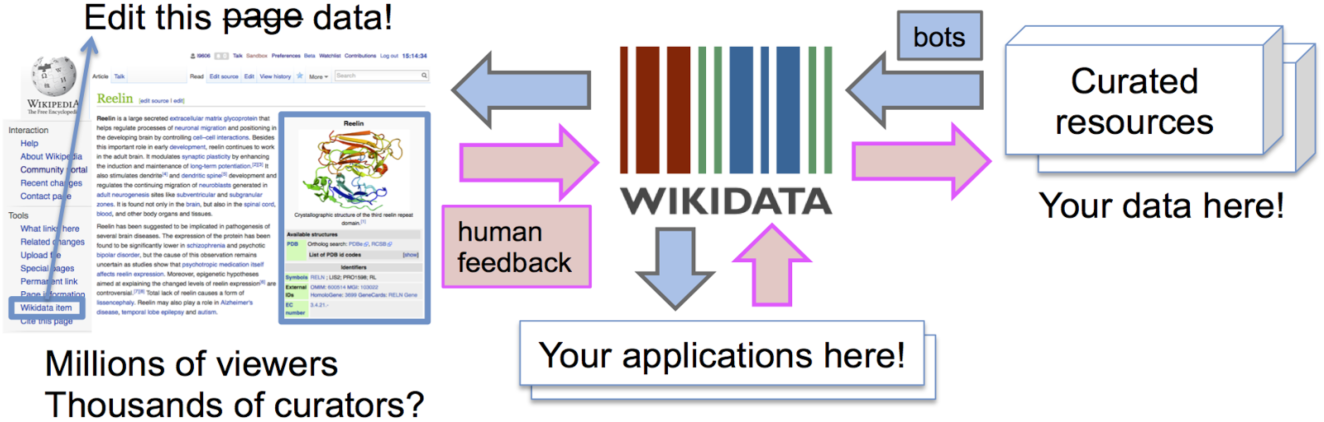
Wikidata provides a centralized resource for structured data. Applications including, but not limited to, Wikipedia can now read and write to Wikidata.

Infoboxes provide the bridge between machine-readable structured data and the unstructured text that forms the main body of each article. Since 2008, the Gene Wiki project has automatically created and maintained the infoboxes for around 10000 articles about human genes [5]. Now, this initiative is focused on generating a foundation of biomedical knowledge in Wikidata that will be used to improve infobox content on Wikipedia and help drive new applications. To date, we have loaded Wikidata with items about: 56451 human and 73086 mouse genes from NCBI Gene [6], 6562 concepts in the Disease Ontology [7], and 1830 FDA-approved drugs. This initial data load generated Wikidata items for these key biomedical concepts, mapped them to Wikipedia articles and linked them to the corresponding identifiers in authoritative public databases. The identifier-level connections to the source databases ensure that Wikidata content can be easily integrated into the existing Web of biomedical data. Moreover, the provenance of all Wikidata claims can be assessed through inspection of the supporting references. The data is kept up to date by periodically running ‘bots’ that propagate changes from authoritative sources to Wikidata. When conflicts arise from human edits to Wikidata items, these are flagged for manual review. The next phase of the project will stitch these concepts into a richly interconnected semantic network.

## 3 Taking a sip of the data soup – Wikidata and the Semantic Web

The first application to use Wikidata extensively is Wikipedia but this could be the tip of the iceberg. To give a preview of what Wikidata could become, it’s useful to briefly examine its closest ancestor, DBpedia. The DBpedia project mines content from Wikipedia by parsing infoboxes, maps this content to their own ontology, and provides access to this data in the form of a large RDF database available both for bulk download and SPARQL query. While enabling interesting queries on its own, its most important function is as a global linking hub for the Semantic Web [8]. In comparison to DBpedia, Wikidata has a number of advantages. First, it can be edited directly and changes are reflected in real time. Second, it does not require any parsing because all data is managed in a database from the outset. Third, it contains large amounts of content that is not present in Wikipedia, such as items for every mouse gene. Finally, its query API supports not only queries along its asserted knowledge graph, but also along references, qualifiers and even edit histories. These additional capabilities, viewed in light of the success of the DBpedia project, portend a vital future for Wikidata in the context of the Semantic Web.

Within the biomedical domain, useful queries are already possible as a result of the ‘single-pot’ nature of Wikidata. For example, it is possible to use Wikidata’s SPARQL endpoint (https://query.wikidata.org/) to answer questions such as “what clinically relevant drug-drug interactions are known for the drug methadone (CHEMBL651)” [9]. **Importantly, the data used to answer this query came from two groups working completely independently**. Our ‘drug_bot’ bot added the CHEMBL identifiers (as well as many other identifiers) while another bot developed by a team at the Medical University of Vienna added the drug-drug interactions [10]. This happened without any direct coordination between our groups.

This kind of serendipitous, automatic, cross-continental data integration is the primary goal of the Semantic Web, but is not yet commonplace. The key beauty and main challenge of the Semantic Web is its distributed nature. In order for this kind of integration to happen in the absence of a centralized resource like Wikidata, several major hurdles would need to be leaped. First, both teams would need to know enough about the fairly complex stack of semantic technologies to provide their data as RDF through a stable, public SPARQL endpoint. Second, they would have to work with overlapping identifier systems. Third, the would-be consumer of their data would need to discover both of their endpoints and be sophisticated enough with SPARQL to identify and issue the appropriate distributed query. All of this is possible and can work, but it is not easy.

By integrating data in a centralized, single community pot, Wikidata provides a platform that addresses each of these problems. Data providers do not have to set up and maintain their own SPARQL endpoint – a challenge that very few teams have succeeded at doing for any length of time [11]. By virtue of working in the same database, it is far less likely – though not impossible – for independent teams to generate and publish different identifiers, as the first step in working with Wikidata is to query it to see what is already there. Finally, the challenge of finding a relevant endpoint is negated when there is only one. Note that Wikidata can be queried using SPARQL or the Wikidata Query Language [12].

## 4 Many Cooks…

The fact that Wikidata is one centralized, community resource immediately surfaces the challenges incurred in any collaborative ontology development process. In Wikidata, the ‘ontology’ corresponds to its collection of linking properties used to describe items. A new property in Wikidata has to be proposed for community discussion and is only created after a consensus regarding the value of the property and its relation to existing properties has been established. For those used to controlling their own data and data models, this process can feel tedious. But this same fundamental process must be undertaken in any attempt at data integration. The fact that it happens up front, when data is first being loaded, should help to keep the data consistent and reduce the downstream identifier and ontological mapping problems that continue to plague bioinformatics.

Imagine the power of combining the structured data in Wikidata, the high accessibility and dedicated community of Wikipedia and the knowledge of the scientific community. Contemplate further that all of this data is freely available and accessible through a stable query interface and robust, read/write API. This makes important, high-quality information easily accessible by anyone and opens up scientific knowledge for public scrutiny. Further, the built-in provenance tracking can provide detailed chains of evidence to support or refute each claim and all of this can be discussed using the many social tools, such as ‘talk pages’ for every data item, baked into the MediaWiki infrastructure.

Aside from creating useful ways to disseminate data, this sociotechnical structure provides a framework for the broad community to broadcast feedback back to the original data owners. Even at this early stage of this project, this process has already led to improvements in source data. For example, in the Disease Ontology the term ‘Ollier disease’ had the synonym ‘Maffucci syndrome’. Upon importing the Disease Ontology into Wikidata, members of the Wikidata community pointed out that the two terms, though putative synonyms, linked to two different extant Wikidata items. Upon closer review it was determined that these two terms represent two different, albeit closely related, diseases, leading to the creation of a new term in the Disease Ontology. As Wikidata expands it is to be expected that additional differences in representation between it and other knowledge resources will surface. These will first be triaged by the WikiData community to check for errors and, if consensus is achieved that there is an error in the original source, this will be relayed for consideration. In this way, the WikiData community can become the ‘many eyes’ that make all ontology bugs shallow.

## 5 …Can Make a Delicious Soup

We can create a powerful commons of biomedical knowledge by building on established resources and the dedicated community to connect genes, proteins, drugs, diseases, phenotypes and symptoms. Wikipedia will be the first application to use the content in Wikidata, but certainly not the last. The fire is ready and the pot is starting to heat up. Some villagers are already peeking out of their windows ready to join us around the pot, but it will take the effort of the whole community to make a delicious biomedical data soup. We invite you to join us in this effort.

